# Contribution of individual excitatory synapses on dendritic spines to electrical signalling

**DOI:** 10.1101/2022.02.15.480607

**Authors:** Ju-Yun Weng, Cesar Ceballos, Dejan Zecevic

## Abstract

Dendritic spines, small (∼1 µm) membrane protrusions from neuronal dendrites which receive most of the excitatory synaptic inputs in the mammalian brain, are widely considered the elementary computational units of the brain. Our understanding of electrical signalling in spines is currently being debated, primarily for methodological reasons. We combined the standard techniques of whole-cell recording and voltage imaging methods to study excitatory postsynaptic potentials evoked by two-photon glutamate uncaging (uEPSPs) on individual dendritic spines on basal dendrites in rat cortical slices. We analyzed the initiation, temporal summation, and propagation of uEPSPs from the spine head to the parent dendrites in three principal neocortical pyramidal neuron classes. Our measurements show no significant attenuation of uEPSPs across the spine neck in most tested mushroom spines on basal dendrites. This result implies that spine synapses are not electrically isolated from parent dendrites and that these spines do not serve a meaningful electrical role. Using the same high-sensitivity voltage imaging techniques, we characterized the temporal summation of uEPSPs induced by repetitive glutamate uncaging mimicking burst activity of presynaptic neurons. We found that responses to high-frequency repetitive quantal EPSPs are strictly limited in amplitude and waveform. This finding reveals a biophysical mechanism for preventing synaptic saturation.

**Significance Statement:** We used an electrochromic voltage-sensitive dye, which acts as a transmembrane optical voltmeter, to define the electrical role of dendritic spines, small membrane protrusions that receive most of the excitatory synaptic inputs in the brain. The data argue that investigated spine synapses of principal neurons are not electrically isolated from the parent dendrites. We also found that the amplitude of temporal uEPSP summation during repetitive synaptic activation is restricted at the site of origin, preventing synaptic saturation. These results facilitate our understanding of how a complex assembly of receptors and ion channels in spines generates and processes electrical signals and mediate plasticity in response to the quantal release of chemical transmitters caused by patterned activity in presynaptic axons.

## Introduction

This study aimed to define the electrical role of dendritic spines and characterize the contribution of single spine synapses to the electrical signalling of individual neurons. Several studies hypothesized that dendritic spines serve a unique electrical role because spine head synapses are electrically isolated from the parent dendrite by the slender spine neck. Other studies concluded that spine synapses are not electrically isolated and that spines have no electrical role. At present, neither one of these opposing postulates is universally accepted. The hypothetical electrical role of spines is a critical issue because it implies high electrical resistance of the ∼1µm long spine neck cable (R_neck_) relative to the input impedance (Z_dendrite_) of the ∼ 60-1000 µm long parent dendritic cables. A high value of R_neck_ would, in turn, imply the functional significance of the highly variable morphology of individual spine necks, making practically every spine functionally different.

One group of studies supporting the electrical role of spines postulate high R_neck_ partially or entirely on theoretical grounds using numerical simulations (Koch et al., 1983; Bloodgood and Sabatini, 2005; Grunditz et al., 2008; Bloodgood et al., 2009; Gulledge et al., 2012; Xu et al., 2012; Araya et al., 2014; Tønnesen et al., 2014; Acker et al., 2016; Cartailler et al., 2018; Lagache et al., 2019). Another group of studies postulates high R_neck_ based on indirect measurements of Ca^2+^-signals from which they derived membrane potential transients. A different indirect measurement approach was to determine diffusional resistance experimentally and derive electrical resistance (Bloodgood and Sabatini, 2005; Araya et al., 2006b, 2006a, 2014; Grunditz et al., 2008; Bloodgood et al., 2009; Harnett et al., 2012; Tønnesen et al., 2014; Bywalez et al., 2015; Acker et al., 2016; Hage et al., 2016; Beaulieu-Laroche and Harnett, 2018). Finally, a third group of studies postulates high R_neck_ based on an attempt to directly probe spine membrane potential changes using electrical and optical techniques (Jayant et al., 2016; Kwon et al., 2017; Cartailler et al., 2018; Cornejo et al., 2022).

In contrast, a different set of reports provides indirect evidence that, in most spines (>80%), the neck resistance is too small relative to the input impedance of the dendrite to affect synaptic signals. Some of these studies are based on theoretical considerations and numerical simulations (Rall and Rinzel, 1973; Rall, 1974; Koch and Zador, 1993). Other studies in this group are based on experimental measurements of diffusional resistance of the spine neck, which indicated relatively low spine neck resistance in the majority of spines (Svoboda et al., 1996; Takasaki and Sabatini, 2014; Tønnesen et al., 2014; Miyazaki and Ross, 2017, 2022). Previously, we provided direct evidence for low electrical resistance of the spine neck as recorded from individual spines on basal dendrites in one class of principal pyramidal neurons. The data were acquired using voltage-sensitive dye recordings with adequate sensitivity and spatiotemporal resolution (Popovic et al., 2015a). This technique uses an organic voltage-sensitive dye that acts as a transmembrane voltmeter with a linear scale in the physiological range of neuronal membrane potential signals. The traces showing fluorescent light intensity changes from the voltage-sensitive probe track the membrane potential transients exactly. The spatial resolution of this method allowed monitoring of uEPSP voltage transients simultaneously at both ends of the spine neck, i.e., in the spine head and the parent dendrite at the base of the spine. In this study, we further improved the temporal resolution of voltage imaging, confirmed earlier conclusions by additional measurements from L5 pyramidal neurons, and extended the experiments to the spines of two other classes of principal pyramidal cells (L2/3 and L6). Under described recording conditions, our experimental results did not depend on assumptions. The obtained data argue for a minimal or no impact of spine necks on electrical signaling in most cases of sampled mushroom spines on basal dendrites of all three classes of neurons. Using the same technique, we monitored the local effects of repetitive activation of individual excitatory synapses and revealed the biophysical mechanism which minimizes synaptic saturation.

## Materials and Methods

### Experimental design and statistical analysis

We used high-sensitivity voltage imaging with an organic electrochromic dye to analyze the electrical role of dendritic spines in selectively labeled L2/3, L5, and L6 pyramidal neurons in rat cortical slices. Statistical analyses were performed using GraphPad Prism 9.3.1. All values are reported as mean ± SEM.

### Slices, patch-clamp recording, and intracellular application of dyes

All surgical and experimental procedures followed the Public Health Service Policy on Humane Care and Use of Laboratory Animals and were approved by Yale University Institutional Animal Care and Use Committee. Experiments were carried out on somatosensory cortex slices from 18-30-day old rats of either sex. The animals were decapitated following deep isoflurane anesthesia, the brain was quickly removed, and 300 μm thick coronal cortical slices were cut in an ice-cold solution using a custom-made rotary slicer with a circular blade (Specialty Blades Inc., Staunton, VA).

Slices were incubated at 34^0^ C for ∼30 minutes and then maintained at room temperature (23-25^0^ C). The standard extracellular solution used during recording contained (in mM): 125 NaCl, 25 NaHCO_3_, 20 glucose, 2.5 KCl, 1.25 NaH_2_PO_4_, 2 CaCl_2_, and 1 MgCl_2_, pH 7.4 when bubbled with a 5% CO_2_ gas mixture balanced with 95% O_2_. Somatic whole-cell recordings in the current clamp or voltage-clamp mode were made with 4-6 MΩ patch pipettes using a Multiclamp 700B amplifier (Axon Instruments Inc., Union City, CA). Voltage-clamp recordings were made with series resistance compensation set at 70%. The pipette solution contained (in mM): 120 K-gluconate, 3 KCl, 7 NaCl, 4 Mg-ATP, 0.3 Na-GTP, 20 HEPES, and 14 Tris-phosphocreatine (pH 7.3, adjusted with KOH) and 0.8 mM of the voltage-sensitive dye JPW3028 (Antić and Zecević, 1995). The pharmacological agents were obtained from Tocris. The somatic whole-cell recording data were not corrected for liquid junction potential. We selected pyramidal cells with intact dendrites in one plane of focus close to the surface of the slice (to minimize light scattering) using infrared differential-interference contrast (DIC) video-microscopy. The recordings were from mushroom spines on superficial basal dendrites at different distances from the soma (60 – 230 µm). Stubby spines without clearly defined spine necks were excluded. This study was restricted to basal dendrites because bAPs used for normalizing the sensitivity of optical recordings from different locations do not propagate into all parts of the apical dendritic arbour. Individual pyramidal neurons were labeled with the membrane impermeant voltage-sensitive dye by allowing free diffusion of the probe from the somatic patch pipette in the whole-cell configuration. We used a voltage probe for intracellular application, JPW3028, synthesized and provided by Leslie Lowe, Centre for Cell Analysis and Modelling, UConn Health Centre. This die is a close analogue of JPW1114 (Zecević, 1996) with similar voltage sensitivity available from Invitrogen as D6923. Glass pipettes were first filled from the tip with the dye-free solution by applying negative pressure for about 15 seconds and then back-filled with the solution containing the indicator dye (0.8 mM). Intracellular staining was accomplished in 15-60 minutes, depending on electrode access resistance. After enough dye diffused into the cell body, as determined by measuring resting fluorescence intensity from the soma, the patch-electrode was detached from the neuron by forming an outside-out patch. The staining level was determined empirically as a compromise that attains an adequate level of fluorescence without causing damage by prolonged dialysis from the patch pipette. The preparation was typically incubated for an additional 1.5 -2 hours at room temperature to allow the voltage-sensitive dye to spread and equilibrate in the dendritic arbour. In order to obtain electrical recordings, the cell body was re-patched using an electrode filled with the dye-free intracellular solution before making optical measurements at 34 C°. Both APs and steady-state hyperpolarizing signals were evoked by transmembrane current pulses delivered via the recording electrode attached to the soma in whole-cell configuration (Popovic et al., 2012).

### Optical recording

The recording setup was built around a stationary upright microscope (Olympus BX51; Olympus Inc., USA) equipped with a high spatial resolution CCD camera for infrared DIC video microscopy (CCD-300-RC, Dage-MTI, Michigan City, IN, USA) and a high-speed data acquisition camera used for voltage imaging. This camera (NeuroCCD-SM, RedShirtImaging LLC, Decatur, GA, USA) is characterized by relatively low spatial resolution (80×80 pixels), exceptionally low read noise, and a full frame rate of 2 kHz. The frame rate can be increased to 5 kHz by reading out the central subsection of the camera chip of 24×80 pixels. The 5 kHz recording mode was used in all experiments to reconstruct signals accurately for calibration and comparison. The brain slice was placed on the microscope stage. A water-dipping objective projected the stained neuron’s fluorescent image onto the CCD positioned in the primary image plane of the microscope. We used a 100X/1.0 NA Olympus objective for optical recordings from individual spines. This objective was a compromise between imaging area, spatial resolution, and signal-to-noise ratio (S/N). The optical recording was carried out in the wide-field epifluorescence microscopy mode. A frequency-doubled 500 mW diode-pumped Nd: YVO4 continuous wave (CW) laser emitting at 532 nm (MLL532, Changchun New Industries Optoelectronics Tech. Co., Ltd., Changchun, China) was the source of excitation light. The laser beam was directed to a light guide coupled to the microscope via a single-port epifluorescence condenser (TILL Photonics GmbH, Gräfelfing, Germany) designed to provide approximately uniform illumination of the object plane. The laser was used as a light source in place of a conventional Xenon arc-lamp to increase the sensitivity of Vm-imaging by (1) providing a monochromatic excitation light at the red edge of the absorption spectrum to maximize the Vm sensitivity of the dye (Loew, 1982; Kuhn et al., 2004; Holthoff et al., 2010; Popovic et al., 2012, 2015a) and (2) increasing the intensity of the excitation light beyond the level that an arc-lamp can achieve. Excitation light was reflected to the preparation by a dichroic mirror with a central wavelength of 560 nm. The fluorescence light was passed through a band pass emission filter (FF01-720/SP-25; 720 nm blocking edge BrightLine multiphoton short-pass emission filter, Semrock). The laser light was gated for voltage imaging by a high-speed shutter (Uniblitz LS6, driver D880C). Acquisition and analysis of data were carried out using NeuroPlex software (RedShirtImaging). In this configuration, a CCD frame (80 × 80 pixels) corresponded to a field of 18 × 18 µm in the object plane, with each pixel receiving light from an area of ∼0.23 × 0.23 µm in the focal plane.

### Computer-generated holography for two-photon uncaging of glutamate

The voltage imaging setup was integrated with an ultra-fast pulsed titanium-sapphire laser tuned to 720 nm for 2-photon glutamate uncaging (Chameleon Ultra, Coherent Inc.). The light intensity of the laser and the duration of the uncaging pulse were controlled by a Pockells cell (Model 350-80, Conoptics Inc.). The two-photon uncaging pattern was generated using a commercial module for holographic illumination (Phasor–3i Intelligent Imaging Innovation, Inc. Denver, CO USA), modulating 720 nm laser source controlled by SlideBook software (3i Intelligent Imaging Innovation). We used multipoint patterns acquired on the NeuroCCD camera to calibrate the exact positioning of the holographic spots. It was possible to achieve submicron precision in spot positioning by introducing a correcting stretch, translation, and rotation transformation to the input patterns provided to the Phasor algorithm. The size of the two-photon 720 nm uncaging spot was measured at the focal plane of the microscope objective illuminated by the beam of parallel light overfilling its back opening. The light was focused on the thin film of rhodamine 6G spin-coated on a coverslip. The induced fluorescence spot was projected onto a CCD camera chip positioned in the primary image plane of the microscope, and its size was measured to be ∼0.6 μm in diameter (Fig. S1). The exact focal volume of 2-photon excitation (Matsuzaki et al., 2001) was not determined, but the spatial resolution of glutamate uncaging was sufficient to activate one spine synapse in isolation. This spatial resolution has been repeatedly documented (Matsuzaki et al., 2001; Smith et al., 2003; Carter and Sabatini, 2004; Sobczyk et al., 2005; Tazerart et al., 2020) and illustrated here in Figure 2. The uncaging 720 nm red light is outside the absorption spectrum of the fluorescent styryl dye JPW302. Nevertheless, we did find that the scattered 720 nm uncaging red light from the focal point caused the Stimulated Emission Depletion (STED) effect reducing the intensity of the voltage-sensitive die fluorescence. However, because we positioned the uncaging light spot at about 0.5 µm away from the spine, because the duration of the uncaging pulse was very short (0.2-0.4 ms), and because the STED effect is instantaneous on the biological time scale, it was possible to vary the position of the uncaging spot around the spine head until only one or sometimes two data points were effected (Fig. S2). Thus, the STED effect did not alter the uEPSP shape and peak amplitude. Wide field illumination was used to obtain an image of dye-loaded dendrites and identify structures of interest for glutamate uncaging and voltage imaging. DNI-glutamate TFA provided by Femtonics KFT (Budapest, Hungary), which has ∼7 times higher 2-photon uncaging efficiency (Chiovini et al., 2014) than the commonly used MNI caged compound, was bath applied at a concentration of 4 mM. The illumination spots were placed at a distance of ∼ 0.5 µm from individual spine heads. The precise spatial relationship between the uncaging spots and spine heads was uncertain at the sub-micrometer spatial scale because of light scattering in the brain tissue. The uncaging light pulse was adjusted in duration (from 0.2-0.5 ms) and intensity (from 10-20 mW under the objective) to produce a response similar to a unitary EPSP in the soma (0.2-0.8 mV) (Magee and Cook, 2000; Nevian et al., 2007; Bloodgood et al., 2009; Enoki et al., 2009).

**Figure 1.**
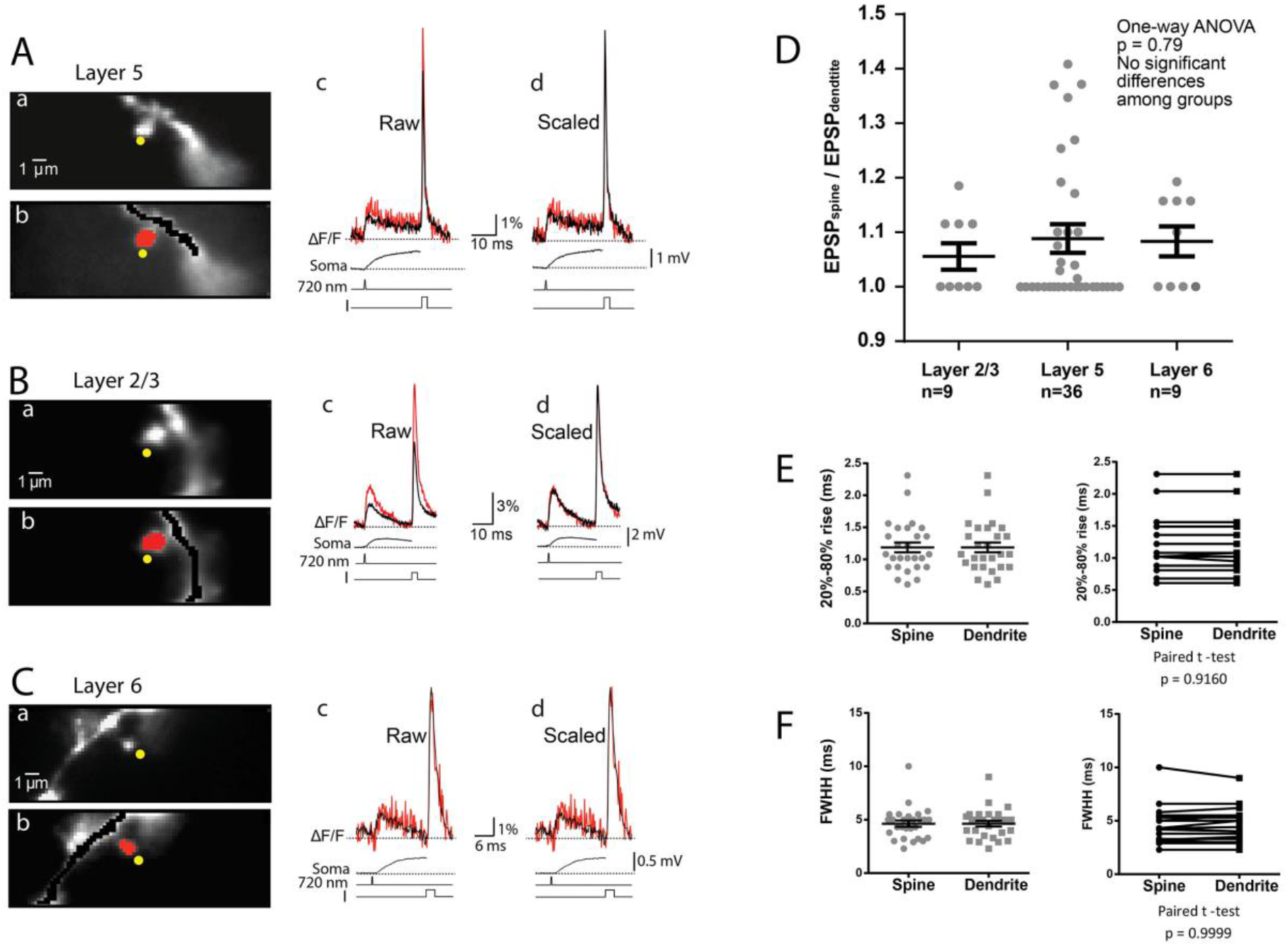
Optical recordings of EPSP and AP signals from individual spines and parent dendrites from cortical L5, L2/3, and L6 pyramidal neurons. **A**, L5 pyramidal neuron. **(a)** Fluorescence image of a spine in recording position obtained with the CCD for voltage imaging. The yellow dots indicate the position of the 720 nm light spot ∼0.6 μm in diameter, used for 2-photon glutamate uncaging. **(b)** Selection of pixels used for the spatial average of optical signals from the spine head (red) and parent dendrite (black). **(c)** Evoked uEPSP and AP signal from spine head and parent dendrite superimposed. Bottom three black traces: Top: electrode recording of somatic uEPSP. Middle: the uncaging command pulse. Bottom: transmembrane current pulse delivered by a somatic patch electrode. **(d)** Superimposed signals from the spine head and parent dendrite corrected for recording sensitivity difference by normalizing recordings to the bAP optical signal. **B** and **C**, Similar recordings from L2/3 and L6 neurons, respectively. **D**, Scatter plot of individual values of the ratio (uEPSPspine/uEPSPdendrite) for the three classes of cortical pyramidal neurons. Vertical lines show mean ± SEM). **E, F**, Scatter plot of individual values of uEPSP 20-80% rise time (E) and of uEPSP full width at half height (FWHH) (F). Vertical lines show mean ± SEM.

**Figure 2.**
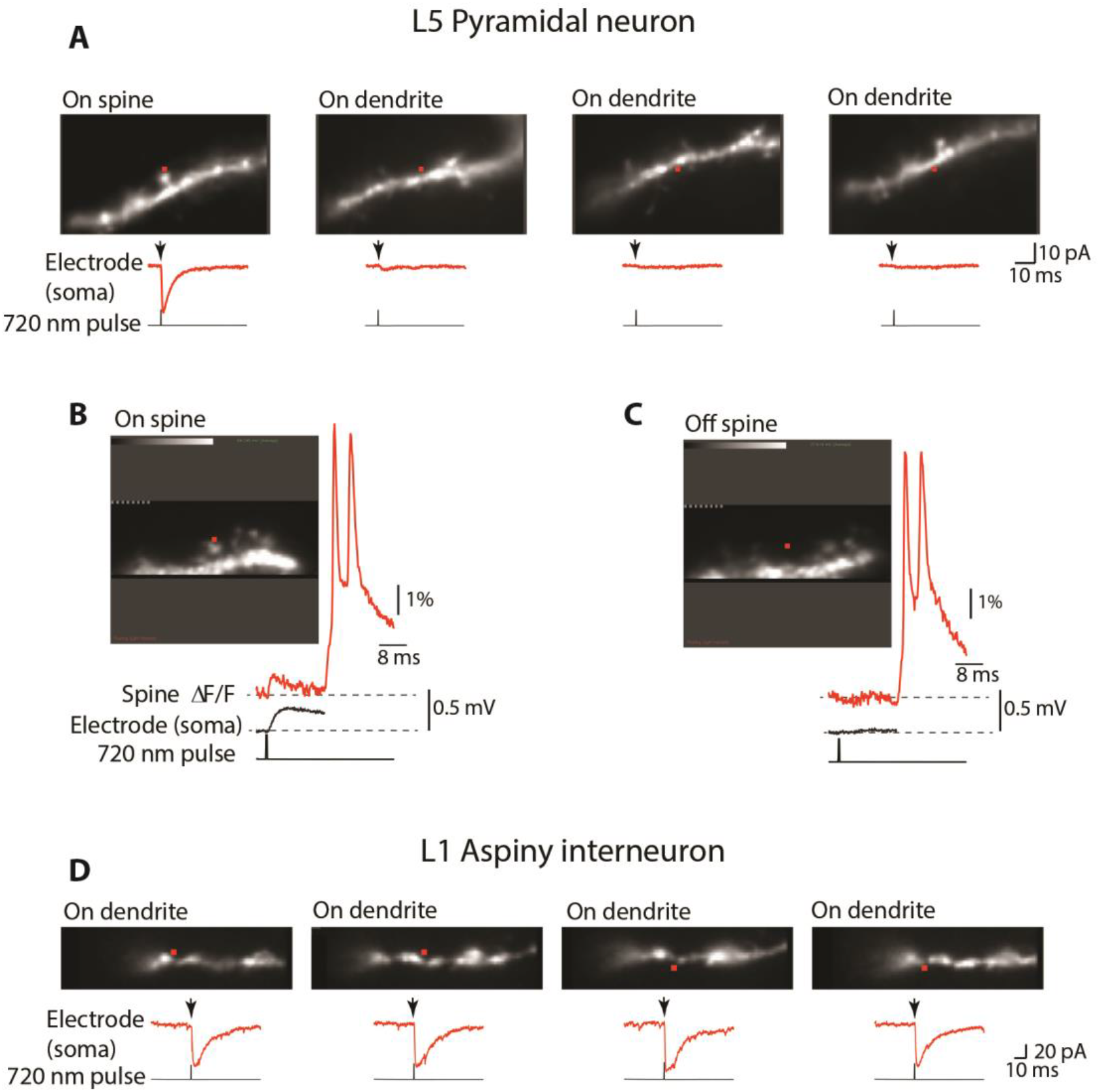
Selective activation of individual synapses. **A**, Uncaging glutamate on smooth, aspiny regions of basal dendrites of L5 pyramidal neurons (On dendrite) evoked currents (red traces recorded under voltage-clamp) that were either not detectable or represented a tiny fraction of the current evoked by activating synapses on neighboring spines (On spine). Red dots indicate the position of the 720 nm light spot used for 2-photon glutamate uncaging. **B**, Uncaging glutamate on an individual spine evokes clear uEPSP as revealed by voltage imaging (red trace) and electrical recording from the soma (black trace). **C**, Uncaging glutamate at a distance from aspiny dendritic regions equivalent to the distance of the spine head failed to produce measurable uEPSP responses in both optical and electrical recordings. **D**, Uncaging glutamate on aspiny dendrites on L1 interneurons evoked a fast and clear response at all tested locations.

At the end of each experiment, a detailed morphological reconstruction of dye-loaded neurons was carried out on a stationary upright microscope (AxioExaminer D1 with zoom tube (0.5 – 4x), Carl Zeiss Microscopy LLC) equipped with a high spatial resolution CCD camera (1392×1024 pixels; Pixelfly-qe, PCO Imaging, Kelheim, Germany) mounted on a spinning-disc confocal scanner (Yokogawa CSU-10). At the end of every experiment, this system collected z-stacks of confocal images for the detailed morphological reconstruction of basal dendrites and spines. Confocal morphological reconstruction was used to confirm that (a) the recorded spine was spatially isolated by more than 10 m from all other spines in both x-y and z-dimensions and (b) to determine the distance of the recording site from the soma. In addition, the recorded length of the basal dendrite was used to calibrate the optical signal in terms of membrane potential using the known bAP amplitude at the corresponding distances (see below).

### Data analysis

Membrane potential optical signals related to (uEPSP) followed by bAPs were recorded typically for 60 ms at a frame rate of 5 kHz at a near physiological temperature of 32-34 C^0^. Data were analyzed and displayed using the NeuroPlex program (RedShirtImaging) written in IDL (Exelis Visual Information Solutions, Boulder, CO) and custom Visual Basic routines. Background fluorescence can be a significant determinant of ΔF/F signal size. Raw data were first corrected for this effect by subtracting the average background fluorescence intensity determined from an unstained area on the slice. Subsequently, signal alignment software was used to correct temporal jitter in AP initiation and possible small preparation movements during averaging (Popovic et al., 2015a). In the temporal domain, AP signals were aligned by cross-correlation of the electrically recorded APs in each trial to the reference signal acquired at the start of averaging. In the spatial domain, camera images were aligned in two dimensions offline by image cross-correlation to compensate for possible small lateral movements of the preparation (Popovic et al., 2015b). The correct focus of the image in the z-dimension was verified before each trial; minor adjustments were often necessary. The spatially and temporally aligned signals were averaged, and slow changes in light intensity due to the bleaching of the dye were corrected by dividing the data by an appropriate dual exponential function derived from the recording trials with no stimulation (Grinvald et al., 1982). The residual slow changes in baseline after bleaching correction, if present, had no effect on uEPSP and bAP amplitude and waveform because they were small and approximately 100 times slower than the rising phase of an action potential (Fig. S3) (Foust et al., 2010; Holthoff et al., 2010; Popovic et al., 2011, 2015a). The waveform of the AP signal was reconstructed from a set of data points using Cubic Spline Interpolation, a piecewise continuous curve passing through each data point (Popovic et al., 2011). Subthreshold optical signals were calibrated on an absolute scale (in mV) by normalizing to an optical signal from a back-propagating action potential (bAP) which has a known declining amplitude along basal dendrites, as previously determined by patch-pipette recordings (Nevian et al., 2007; Palmer and Stuart, 2009). These methods of calibration produce the same results as normalizing signals to optical recordings corresponding to long hyperpolarizing pulses delivered to the soma, which attenuate relatively little as they propagate along dendrites (Stuart and Spruston, 1998; Nevian et al., 2007; Palmer and Stuart, 2009; Holthoff et al., 2010).

## Results

### Majority of spine synapses are not electrically isolated from basal dendrites

The first series of experiments aimed to determine whether our previous findings in L5 cortical neurons (Popovic et al., 2015a) are valid for other principal cortical pyramidal cell classes. The experiments were conducted on basal dendrites of L2/3, L5, and L6 pyramidal neurons. The only direct way to determine the degree of electrical isolation of spine synapses is to simultaneously record EPSP signals from the spine head and the parent dendrite at the base of the spine following selective quantal activation of a single synapse. Under these conditions, the evoked EPSP signal in the spine head represents a voltage drop caused by the synaptic current flow across R_neck_ and Z_dendrite_ connected in series according to the expression:

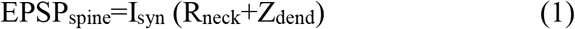

The EPSP signal in the parent dendrite at the base of the spine represents a voltage drop caused by the same synaptic current flow across Z_dendrite_ alone, according to the expression:

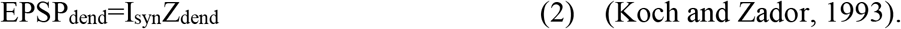

According to these two expressions, the experimental approach to the electrical role of spines is conceptually trivial, even though it is technically demanding to the degree that prevented these measurements for several decades. If one can simultaneously record EPSP_spine_ and EPSP_dendrite_, at the required spatiotemporal resolution, one can anticipate two different categories of results. One possible result is that the EPSP amplitude is significantly larger in the spine head than in the parent dendrite. This result would imply a relatively large R_neck_ comparable to Z_dendrite_ at a given dendritic location. For example, if the amplitude of the recorded EPSP_spine_ is two times larger than EPSP_dendrite_, which would mean that R_neck_ is equal to Z_dendrite_. The other possible result is that EPSP_spine_ and EPSP_dendrite_ are not significantly different in size and shape. A direct conclusion from this type of result would be that R_neck_ must be negligible compared to Z_dendrite_.

We used 2-photon uncaging of glutamate and high-sensitivity organic voltage-sensitive dye recordings to monitor simultaneously quantal uEPSP signals from individual dendritic spine heads and parent dendrites at the base of the spine in three classes of principal pyramidal neurons in L2/3, L5, and L6 of the somatosensory cortical slices. In order to minimize photodynamic damage, the duration of recordings was kept to a minimum, necessary to include the uEPSP and bAP signal within the same recording period. When bAPs were slow, the full repolarization phase was not captured at dendritic locations far from the soma. Recordings were carried out at a frame rate of 5 kHz (using a central subsection of 24×80 pixels of the NeuroCCD-SM; Methods) to improve the accuracy of our earlier results obtained at the sampling rate of 2 kHz (Popovic et al., 2015a). The results (Fig. 1) documented that the method has adequate sensitivity at the required spatiotemporal resolution to simultaneously record and faithfully reconstruct individual EPSPs and APs signals from spine heads and parent dendrites at near physiological temperature. It is noteworthy that this is the only methodology to accomplish this goal. Fig. 1A-C illustrates optical recordings of the uEPSP signals adjusted in uncaging light intensity and duration (Methods) to mimic quantal glutamate release resulting from the arrival of one AP at the presynaptic bouton. Immediately following the uEPSP, within a 60 ms recording period, we evoked a bAP by a brief depolarizing current pulse delivered from the somatic patch electrode.

In all measurements, we used the bAP-related optical signals in spines and dendrites to normalize the sensitivity of optical recordings from different locations (scaled signals). This normalization is based on prior knowledge that bAP has the same size and shape in the spine and the parent dendrite(Palmer and Stuart, 2009; Holthoff et al., 2010; Popovic et al., 2014). The experimental data in Fig.1 show that, in the preponderance of cases, there was very little or no detectable difference in both amplitude and kinetics between EPSP_spine_ and EPSP_dendrite_, a result that implies that R_neck_ is negligible compared to Z_dendrite_. The same result was obtained from L2/3, L5, and L6 pyramidal neuron measurements. Combined with our previous data from L5 pyramidal neurons (Popovic et al., 2015a), the summary plot in Fig. 1D shows that negligible R_neck_ was confirmed in 9 spines from L2/3 neurons, 36 spines from L5 neurons, and 9 spines from L6 neurons. The average EPSP_spine_/EPSP_dendrite_ ratio from all experiments was 1.08±0.03 (n=54) with no significant difference between the three classes of neurons (One-way ANOVA; p=0.79). While the average ratio from 54 experiments was near 1.0, we recorded several outliers with a ratio as high as 1.4. A small percentage (<5%) of spines characterized by high and spontaneously reversible diffusional resistance has been described earlier (Bloodgood and Sabatini, 2005) (see Discussion). The scatter plots showing the distribution of individual values for the kinetics (rise time and FWHH) for EPSP_spine_ and EPSP_dend_ are shown in Fig. 1E-F. The results from basal dendrites of three different classes of principal cortical neurons provide direct evidence that, in the vast majority of mushroom spines, R_neck_ is negligible relative to Z_dendrite_. We conclude that synapses on these spines are not electrically isolated from the parent dendrite to the degree that would imply functional meaning. These findings do not depend on any assumptions when supplemented by the necessary control experiments carried out to address three possible sources of errors that could influence the accuracy of our measurements and the validity of our conclusions: (1) insufficient sensitivity of voltage imaging; (2) contribution of extrasynaptic receptors; (3) insufficient spatial resolution of voltage imaging. The elementary methodological issues in voltage imaging regarding linearity in fractional fluorescence light intensity changes with membrane potential, the absence of pharmacological effects of the JPW3028 dye, the absence of photodynamic damage under correct experimental conditions, as well as the absence of significant effects of the slow bleaching of the dye minimized by correction procedures, have been analyzed in detail and repeatedly documented in both pioneering studies (Ross et al., 1977; Grinvald et al., 1982) and our earlier reports (Djurisic et al., 2004; Canepari et al., 2007; Foust et al., 2010; Holthoff et al., 2010; Popovic et al., 2011, 2015a). In light of the results from a number of these control experiments, it is unlikely that pharmacological effects or photodynamic damage caused by the die specifically affected R_neck_. However, it was not possible to definitely exclude this possibility. This is because an alternative method to provide direct evidence on R_neck_ without a voltage-sensitive die has not been worked out.

### Sensitivity of optical recording

The accuracy of our measurements depends directly on the sensitivity of optical recording at the spatiotemporal resolution required for the correct reconstruction of electrical signals from optical data. Figure 1 indicates that our recordings at the frame rate of 5 kHz and spatial scale of individual spines can resolve both bAP and EPSP signals with a signal-to-noise ratio ranging from 3 to 20 in different experiments. We have documented previously that a sampling rate of 5 kHz (but not 1 or 2 kHz) was adequate for accurate reconstruction of the bAP in both dendrites and spines (see Fig. 4 in Popovic et al. 2014). Accurate reconstruction is critical because AP signals normalize optical recording sensitivity from different locations. It follows that uEPSP signal reconstruction was also accurate because uEPSPs have slower dynamics than bAP signals. Thus, it is safe to conclude that comparing uEPSP from spines and dendrites was accurate after normalizing the recording sensitivity from different regions.

### Contribution of extrasynaptic receptors

The validity of our results depends on the spatial selectivity of the 2-photon uncaging of glutamate. If 2-photon uncaging on spine head synapses also significantly activated extrasynaptic receptors on parent dendrites, this effect would contribute to the recorded similarity of responses from spines and dendrites. Both pioneering and the most recent studies firmly established single spine spatial resolution of 2-photon uncaging (Matsuzaki et al., 2001; Smith et al., 2003; Tazerart et al., 2020). Furthermore, several laboratories investigated and documented that the non-specific activation of glutamate receptors on parent dendrites is so small that it can be safely neglected (Matsuzaki et al., 2001; Sobczyk et al., 2005; Popovic et al., 2015a).

We confirmed this conclusion in voltage-clamp experiments (Fig. 2A), showing that uncaging glutamate on aspiny membrane regions of basal dendrites of pyramidal neurons at the range of light intensities used in our measurements (Methods) evoked currents that were either not detectable or represented a tiny fraction (4%± 0.9%; n=34) of the current evoked by activating synapses on neighboring spines. Moreover, uncaging glutamate at a distance from aspiny dendritic regions of pyramidal neurons equal to the distance of the spine head failed to produce any measurable uEPSP responses (Fig. 2B, C). This result was confirmed in 9 experiments. In contrast to these results on pyramidal neurons, and as a positive control, we found that uncaging glutamate directly onto the membrane of aspiny dendrites on L1 interneurons evoked a fast and clear response at all locations tested in n=18 interneurons (Fig. 2D). This result is in line with previous reports showing that AMPA glutamate receptors are mainly absent from extrasynaptic regions on basal dendrites of cortical pyramidal neurons (Carter and Sabatini, 2004; Sobczyk et al., 2005; Higley et al., 2012; Popovic et al., 2015a) while they are widely distributed along densely innervated smooth dendrites of aspiny interneurons (Gulyás et al., 1999; Sancho and Bloodgood, 2018).

### Light scattering and spatial resolution

Light scattering in the brain tissue limits the spatial resolution of voltage imaging in wide-field microscopy mode. Because the uEPSP in the dendrite is secondary to the uEPSP in the spine and, hence, must always be smaller, the relevant question is whether fluorescent light from spine heads contaminated optical signals from the parent dendrite, making it artificially bigger. This type of crosstalk, if significant, would make the recorded signals from spines and dendrites similar. However, it has been established that the extent of this contamination in the superficial layers of the slice, less than 30 μm below the surface, was minimal and often not detectable (∼ 3%; see Figs. 3 in Popovic et al. 2014; 2015b). Furthermore, the crosstalk in the opposite direction (from the parent dendrite to the spine head) was also shown previously to be small in superficial layers of the slice (<10%; see Figs. 3 in Popovic et al. 2014; 2015b) and without significant effect on our conclusions.

**Figure 3.**
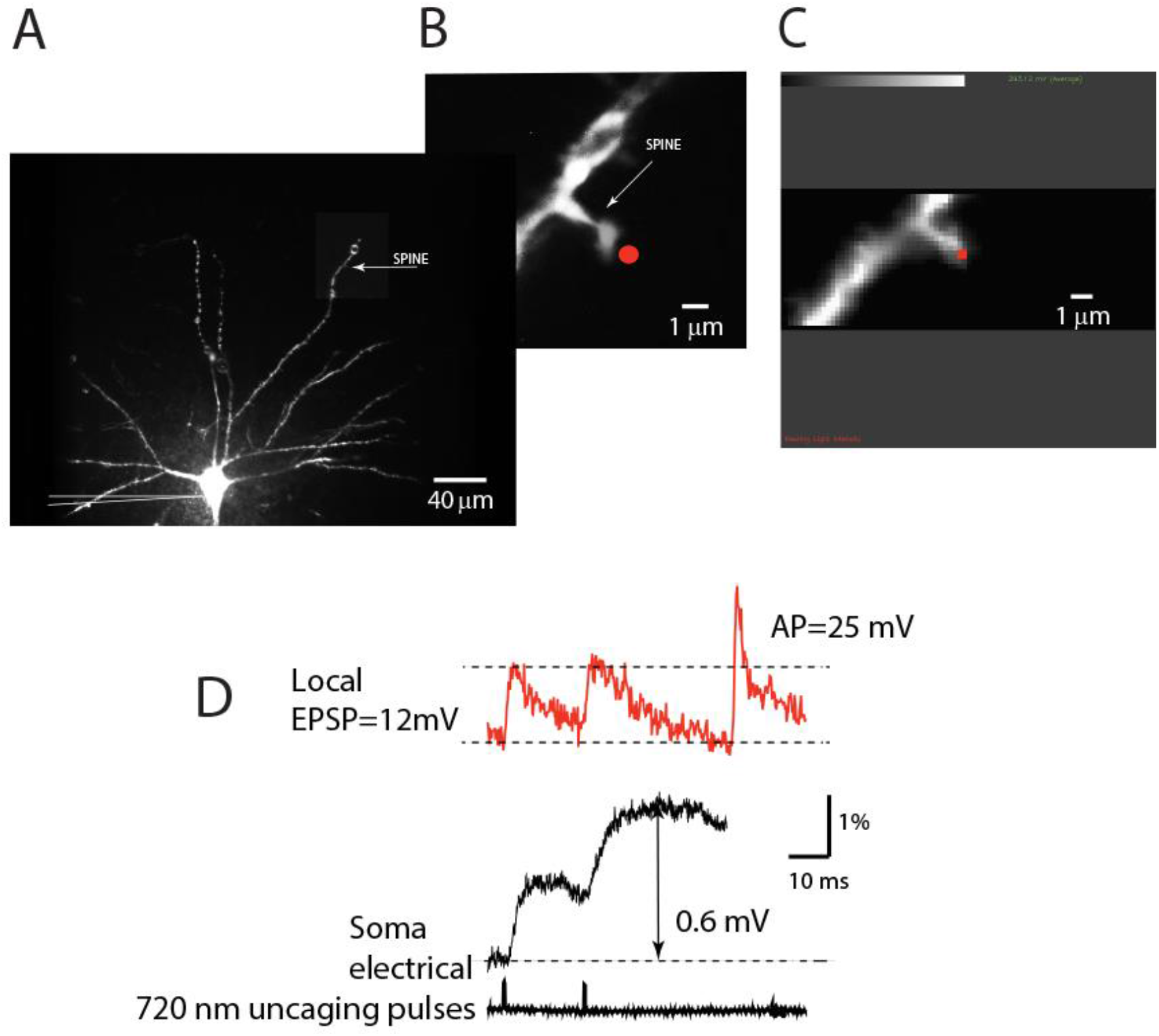
Contribution of individual synapses: experimental design. **A**, An individual neuron is labeled with a voltage-sensitive dye. An isolated spine is identified in a distal region of basal dendrite under low magnification. A patch electrode (shown schematically) is attached to the soma for electrical recording and stimulation. **B**, Fluorescent image of an isolated spine in recording position obtained at high magnification with a high-resolution CCD. **C**, the image of the same spine obtained by reading out a 24×80 pixel subsection of the CCD camera for voltage imaging at 5 kHz. The red dot indicates the position of the 720 nm light spot used for 2-photon glutamate uncaging. **D**, Red traces: optical recordings of local uEPSP signals evoked by 2-photon glutamate uncaging followed by a bAP signal evoked by depolarizing pulse delivered by the somatic patch electrode. Optical signals are spatial averages of all bright pixels in the image. Black traces: somatic electrode recordings (upper) and uncaging command pulses (lower).

**Figure 4.**
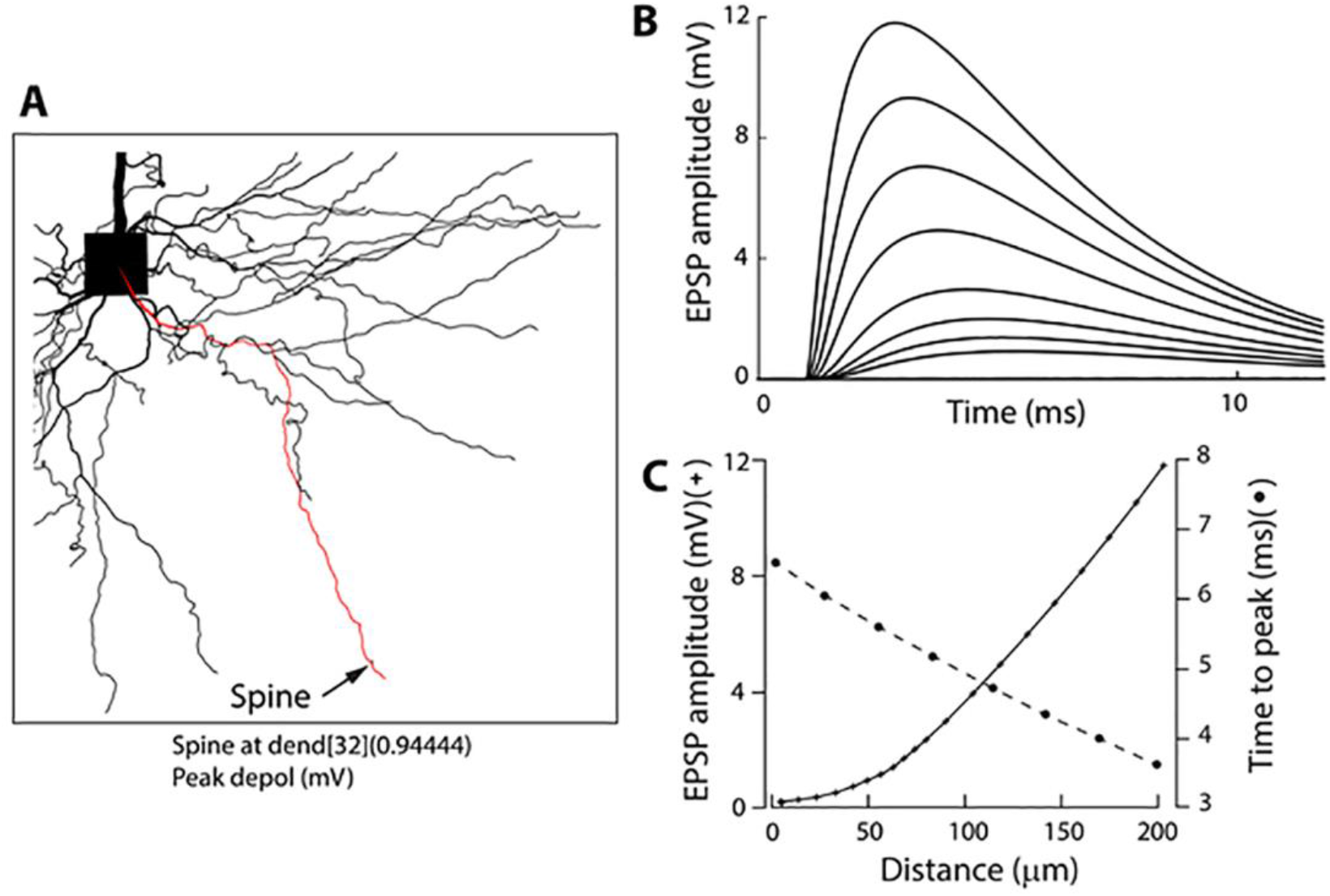
Numerical simulation (NERURON 6.0) of the electrotonic spread of an EPSP along a basal dendrite. **A**, Morphometrically detailed reconstruction of basal dendrites obtained with high-resolution spinning disk confocal microscope. The location of a spine at the distal end of a basal dendrite is indicated by an arrow. **B**, Shape, and size of an EPSP from multiple locations at increasing distances, indicated in (C), from the site of origin (spine) along the basal dendrite. **C**, EPSP amplitude and time-to-peak as a function of distance from the soma.

It is probably helpful to reiterate that the comparison showing similar size and shape of uEPSP signals from spine heads and parent dendrites due to R_neck_<<Z_dendrite_ does not depend on any assumption. The ratio (uEPSPspine/uEPSPdendrite) does not depend on absolute values of R_neck_ and Z_dendrite_ and, therefore, does not require calibration of optical signals in terms of membrane potential. Accordingly, possible errors due to unavoidable inaccuracies in calibrating optical signals in terms of membrane potential (Popovic et al., 2015a; Acker et al., 2016; Kwon et al., 2017; Cornejo et al., 2022) can safely be ruled out. Furthermore, the data argue that uEPSPs are not significantly attenuated as they propagate from synapses on spine heads to the parent dendrite. This result implied that, in electrical terms, synapses on cortical dendritic spines behave in the same way as synapses made directly on dendrites.

### Temporal summation at single spines

The ability to monitor local electrical signaling from individual spines and parent dendrites allowed us to record and analyze temporal summation of uEPSPs following repetitive activation of single spine synapses. Because presynaptic neurons often fire in a burst (Kole, 2011), these experiments mimic physiological conditions in which the natural sensory stimulus activates isolated individual spines on dendritic branches *in vivo* (Jia et al., 2010; Chen et al., 2011; Varga et al., 2011). Figure 3 illustrates the experimental approach for monitoring temporal summation at individual synapses. A fluorescence image of an L5 pyramidal neuron labeled with the voltage-sensitive dye was used to identify an isolated spine close to the surface of the slice (Fig. 3A).

Using patterned illumination based on the CGH system (Methods), a 720 nm light spot from a pulsed laser was positioned within 0.5 μm from the edge of an individual spine for 2-photon uncaging of glutamate. A patch electrode was attached to the cell body to monitor the membrane potential and synaptic currents. The electrode also allowed us to pass depolarizing current and evoke a bAP used to calibrate optical signals on an absolute scale (Methods). The uncaging light pulse was adjusted in duration and intensity to mimic a unitary EPSP in the soma (0.2-0.8 mV; Methods). In order to optimize sensitivity in this set of experiments, uEPSP-related optical signals were recorded at the site of origin as the spatial average of the spine and a small region (∼15 um) of the dendrite at the base of the spine. We showed above (Fig. 1) that this entire surface is very nearly equipotential. Optical signals were recorded simultaneously with the electrode recording from the soma (Fig 3D). From this type of measurement, we established that an average quantal uEPSP recorded at the site of origin had a 20-80% rise time of 1.2 ± 0.07 ms and FWHH of 4.6±0.3 ms (n=26). Scatter plots showing the distribution of individual values are shown in Fig. 1E-F. Due to the rapid kinetics of EPSPs at the site of origin, there was very little or no temporal summation of signals at the synapse if the uncaging pulses were delivered with an inter-pulse interval ≥ 20 ms (Fig 3D). However, a clear summation was recorded at the soma-axon region. This is because EPSP in the soma had considerably reduced amplitude and slower kinetics (Fig. 3D). This result is expected due to the well-known electrotonic propagation and RC filtering effect on electrical signals in the dendritic cable(Rall et al., 1967). We confirmed the declining electrotonic propagation and the RC filtering effect on the dynamics of EPSPs in basal dendrites using numerical simulation (Fig. 4). A detailed treatment of the computational model is available elsewhere (Djurisic et al., 2008; Popovic et al., 2014, 2015a).

In order to ensure that local summation will occur, the following measurements were carried out with 5 uncaging pulses delivered at 200 Hz, mimicking the burst of APs in a presynaptic neuron. At the start of the experiment, we made somatic electrical recordings of uEPSCs under voltage clamp in response to repetitive activation of a spine synapse. Figure 5B illustrates a typical response showing that synaptic currents caused by individual uncaging pulses summated in a pronounced sublinear fashion. A plateau was reached after the second pulse in this experiment, and the maximum current during the plateau phase reached a peak value of 70 pA. The summary data from an extensive series of similar measurements show that the average maximum summated current amplitude during the plateau phase was 52±1.7 pA (n=94). The distribution of individual values is shown in the scatter plot (Fig. 5D).

**Figure 5.**
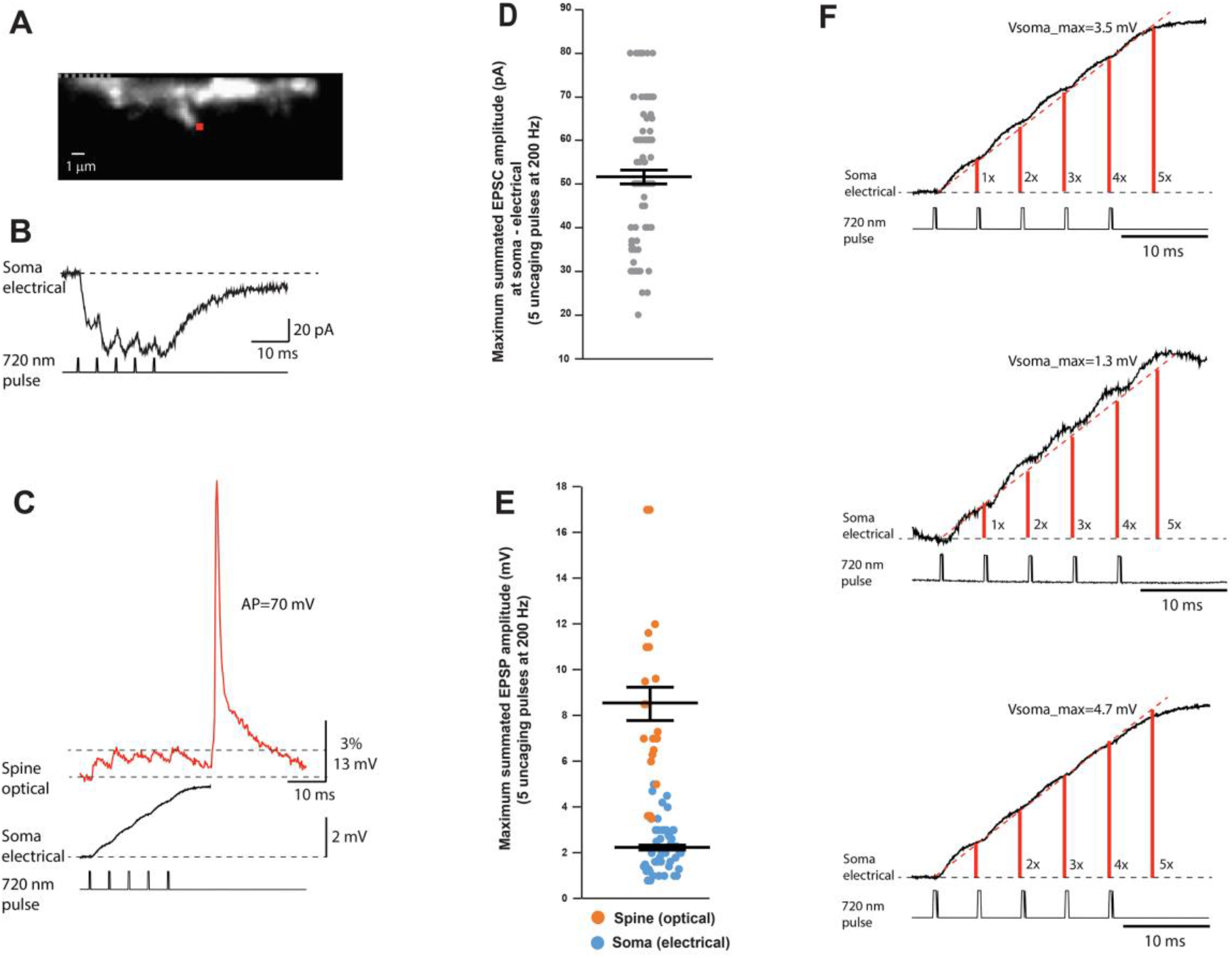
Summation of uEPSP signals. **A**, Fluorescence image of a spine in recording position. A red dot indicates the position of the 720 nm light spot used for 2-photon glutamate uncaging. **B**, Upper trace: summated uEPSCs recorded with a somatic patch electrode under voltage-clamp in response to repetitive glutamate uncaging at 200 Hz. Lower trace: command uncaging pulses. **C**, Upper trace: Summating uEPSP-related signal recorded optically from the dendritic spine and the parent dendrite section shown in (A) in response to repetitive glutamate uncaging at 200 Hz. Middle trace: Summating uEPSP signal recorded with a somatic patch electrode simultaneously with the optical recording of local signals. Bottom trace: Uncaging command pulses. **D**, Scatter plot of individual voltage-clamp measurements of maximum summated uEPSC amplitudes in response to 5 uncaging pulses at 200 Hz. **E**, Scatter plot of maximum summated uEPSP amplitudes in response to 5 uncaging pulses at 200 Hz as recorded optically from the spine and electrically from the soma. **F**, Magnified display of uEPSP trains recorded with a somatic patch electrode (as in middle trace in C) indicating three typical examples of consistent linear summation of subthreshold signals at the soma-axon region.

The voltage-clamp measurement of synaptic current was followed by an optical recording of local uEPSPs in current-clamp mode evoked by an identical uncaging protocol from the same spine on the basal dendrite. Fig. 5C indicates a typical example of a prominent sublinear summation of local uEPSPs consistent with the summation profile of synaptic currents. The maximum response was reached after the second EPSP in this experiment, with the local depolarization reaching a plateau at 6 mV. At the same time, the summation of attenuated uEPSP signals measured at the soma with patch electrode was nearly linear, resulting in the maximum somatic depolarization of 3.5 mV. In a series of measurements of this kind, the average peak amplitude of the summated uEPSP at the site of origin was 8.8±1.0 mV (n=23). The corresponding summated uEPSP peak amplitude in the soma was 2.3±0.2 mV (n=56). The distribution of individual values of the summated uEPSP in the spine and the soma is shown in the scatter plot in Fig. 5E. The RC filtering effect of basal dendrites and the cell body promotes an approximately linear summation at the soma-axon region by slowing the kinetic of EPSP signals as they propagate to the soma. Fig. 5F shows three additional typical examples of linear summation of the train of uEPSPs as recorded in the soma with a patch electrode. An important advantage of this biophysical property of neurons is that it provides a mechanism for a wide dynamic range of near-linear integration of synaptic signals at the soma-axon region (the output end of the neuron) while minimizing saturation of the driving force for synaptic current at remote local synapses on spines (the input end of the neuron).

Because individual synapses are critical computational units in the nervous system, it is important to investigate biophysical properties that determine the sublinear temporal summation of subthreshold signals in dendritic spines. It is clear that relatively small local dendritic depolarization caused by summated uEPSP train (<10mV; Fig. 5E) cannot explain the pronounced sublinear temporal summation based on the reduction of a large driving force for sodium ions (V_Na_=60.60 mV), the dominant current carriers underlying EPSPs (Miyazaki and Ross, 2022). Additionally, glutamate receptors on the postsynaptic side are far from saturation during quantal transmission (Liu et al., 1999; McAllister and Stevens, 2000). Thus, it is likely that AMPA receptor desensitization (Kiskin et al., 1986) plays an important role in the sublinear summation of unitary uEPSPs shown in Fig. 5. To test this prediction, we monitored uEPSCs summation under voltage-clamp as recorded by a patch-electrode at the soma under control conditions and after AMPA receptor desensitization was inhibited by 100 µM cyclothiazide (CTZ) (Partin et al., 1993) (Fig. 6). CTZ is known to produce a fast inhibition of AMPA receptor desensitization (Fucile et al., 2006). On average, adding 100 µM cyclothiazide increased the maximum synaptic current response from 52±2.1 pA to 177±21 pA (n=9), an increase in the mean value of 340%. The scatter plot of the summary data (Fig. 6C) shows the distribution of individual values. The effect was clear and statistically significant, as indicated by paired t-test in Fig. 6D. In cyclothiazide, with desensitization abolished, the synaptic current responses still summate in a sublinear mode. However, the maximum current amplitude level is shifted toward larger values. In these conditions, the plateau is likely caused by receptor saturation. One would expect that almost all synaptic AMPA receptors are activated at saturation, so a further application of glutamate will not increase synaptic current significantly. Indeed, this result was obtained when the train was extended from 5 to 10 uncaging pulses (Fig. 6E). The mean plateau value of the summated EPSC following 10 uncaging evoked releases of glutamate reached a value of 218±36 pA, a slight increase which was not statistically significant (t-test, p=0.48, n=6) (Fig. 5F).

**Figure 6.**
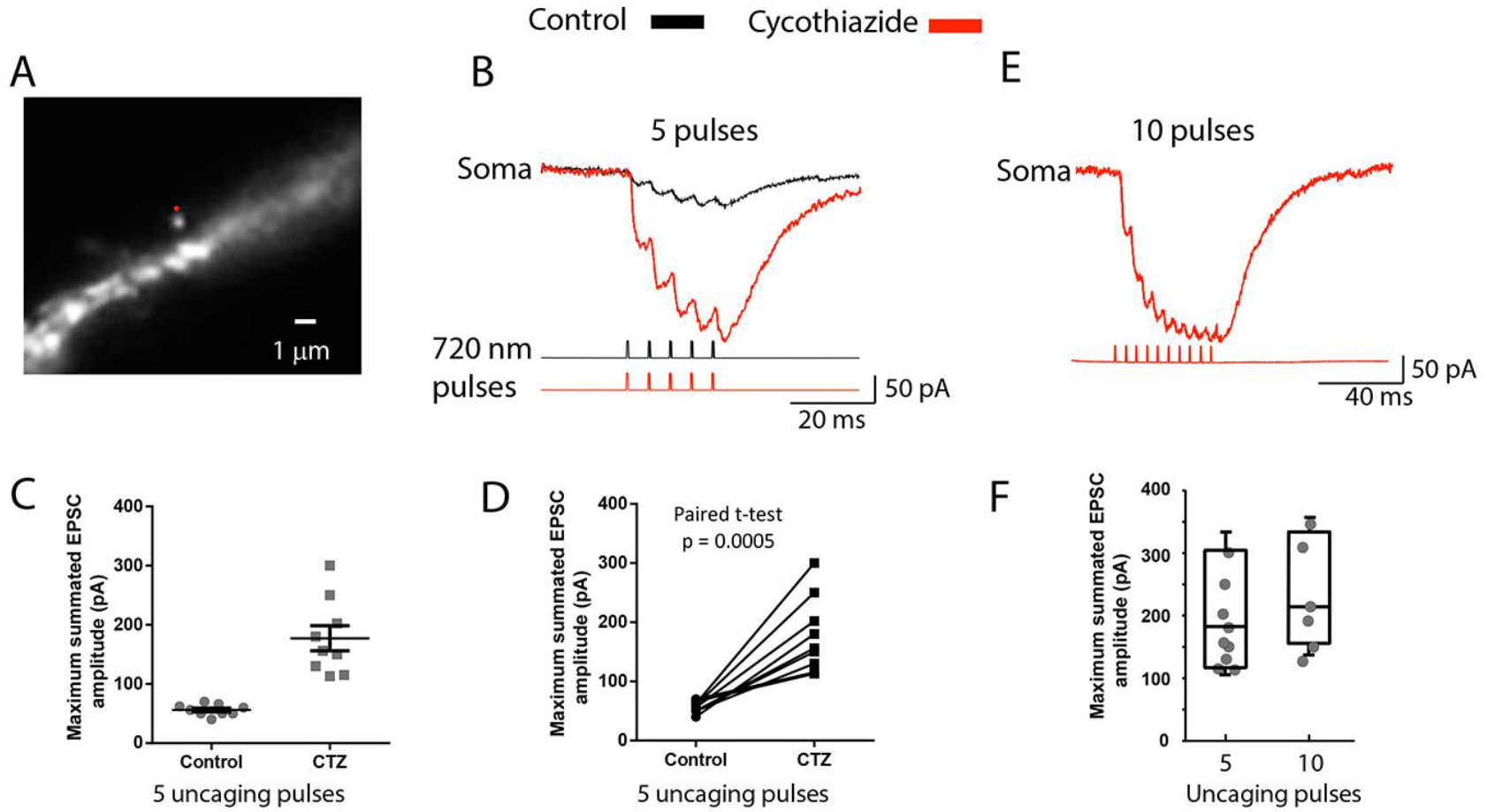
AMPAR desensitization underlies the sublinear summation of unitary uEPSCs. **A**, Fluorescence image of a spine in recording position. Red dot indicates the position of the 720 nm light spot used for glutamate uncaging. **B**, Summating uEPSCs recorded with a somatic patch electrode under voltage-clamp in response to repetitive glutamate uncaging at 200 Hz under control conditions (black traces) and following bath application of 100 µM of AMPAR desensitization inhibitor cyclothiazide (red trace). Cyclothiazide caused a dramatic increase in the maximum synaptic current response. Bottom traces: Uncaging command pulses. **C**, Scatter plot showing the distribution of data from individual experiments. **D**, Paired t-test. **E**, Summating uEPSCs recorded with a somatic patch electrode under voltage-clamp in response to 10 glutamate uncaging pulses at 200 Hz following bath application of cyclothiazide. **F**, Comparison of mean plateau values of the summated EPSC following 5 and 10 uncaging pulses.

Following the somatic EPSC recordings in the voltage-clamp mode, we determined the effects of the desensitization block on the train of local EPSPs measured optically as the spatial average of the activated dendritic spine and a small section of the parent dendrite at the base of the spine (Fig. 7). Because the recordings were carried out from the same location, normalizing the sensitivity was not needed, and the results were directly compared.

**Figure 7.**
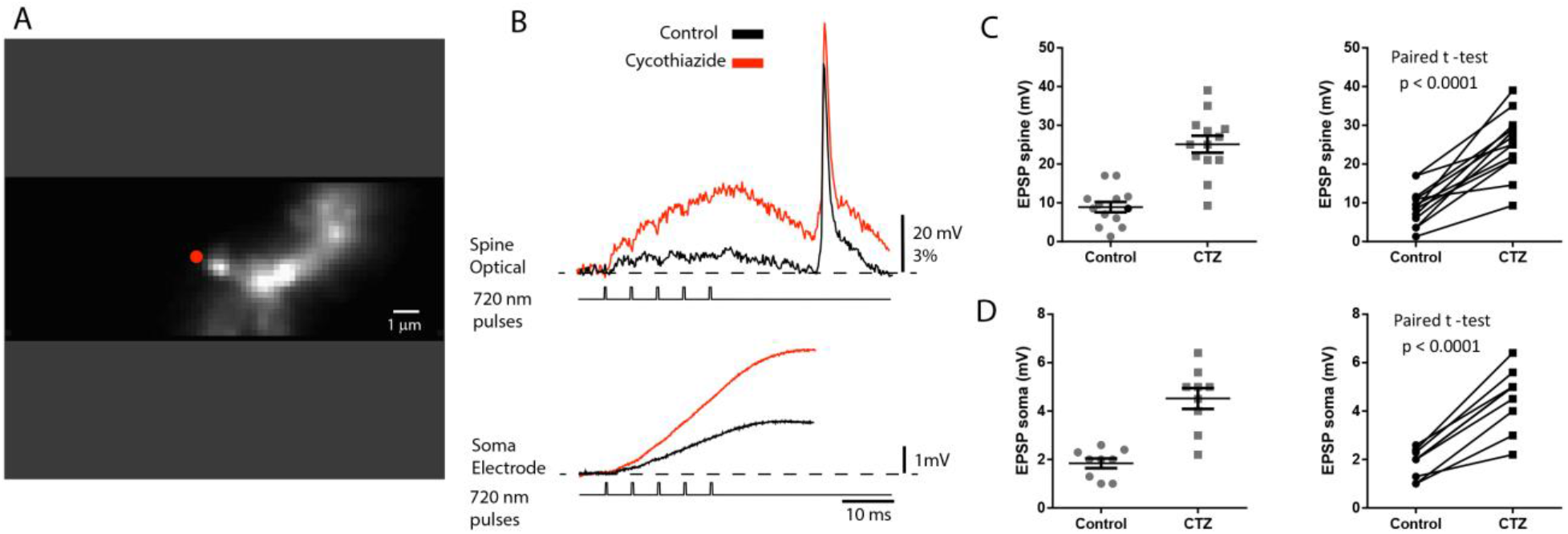
The effect of desensitization on the summated train of uEPSPs. **A**, Fluorescence image of a spine in recording position. The red dot indicates the position of the 720 nm light spot used for 2-photon glutamate uncaging. **B**, Upper traces: optical recordings of local summated uEPSP signals from the spine and parent dendrite under voltage-clamp evoked by repetitive glutamate uncaging at 200 Hz. Black trace: control conditions. Red trace: bath application of 100 µM AMPAR desensitization inhibitor cyclothiazide. Lower traces: simultaneous somatic patch electrode recordings. Bottom traces: uncaging command pulses. **C**, Left panel: scatter plot of individual values of maximum summated uEPSP measured optically in the spine under control conditions and following bath application of cyclothiazide. Right panel: paired t-test shows a significant difference. **D**, Left panel: scatter plot of individual values of maximum summated uEPSP measured with somatic patch electrode under control conditions and following bath application of cyclothiazide. Vertical lines show mean ± SEM. Right panel: paired t-test indicates significant difference.

The local EPSP response measured optically in the spine and a small dendritic section at the bottom of the spine increased from 8.4±0.3 mV in the control solution to 25±3.4 mV (n=13), an increase of the mean value of 297%, following bath application of cyclothiazide. On average, the cyclothiazide caused a 244% increase in summated EPSP response in the soma, from 1.8±0.2 mV to 4.4±0.4 mV (n=9). The scatter plot of the summary data (Fig. 7C-D) shows the distribution of individual data and the high statistical significance of the effect.

## Discussion

### EPSP transfer from the spine to dendrite

The optical recording data show that the electrical resistance of the spine neck in most probed L5 pyramidal neurons is too low, relative to the input impedance of the parent dendrite, to cause a voltage drop that would contribute significantly to the amplitude of the EPSP in the spine head. Similar experiments on L2/3 and L6 pyramidal neurons indicated that the same conclusion is valid for the two additional classes of principal cortical neurons. These results rule out the electrical role of examined mushroom spines on basal dendrites of cortical pyramidal neurons. Our findings are based on measurements of the amplitude ratio (AR) of optical signals AR = EPSP_spine_/EPSP_dend_ = 1+ (R_neck_/Z_dend_) rather than on an attempt to measure absolute values of R_neck_ and Z_dendrite_. The recorded values of the functional parameter AR, which controls the synaptic weight, are, on average, very close to 1, indicating that Rneck is negligible relative to Z dendrite. This finding derived directly from optical recordings does not depend on any assumption. Our measurements also revealed the existence of a small subset of spines with a ratio as high as 1.4 (Fig. 1D). Strong evidence for a small percentage of spines (∼5%) characterized by unstable and spontaneously reversible (on a minutes’ scale) high diffusional isolation of spine heads corresponding to high Rneck and AR has been reported earlier (Bloodgood and Sabatini, 2005). Our study did not examine the stability of high AR over time in these rare cases. The specificity of these spines is presently unclear.

Our results showing relatively low R_neck_ agree with early theoretical predictions (Rall, 1974; Koch and Zador, 1993) and our numerical simulations with a range of plausible biophysical parameters (Popovic et al., 2014, 2015a) as well as with the initial voltage-sensitive dye study of subthreshold signals from dendritic spines carried out using low-sensitivity confocal imaging in combination with numerical simulations (Palmer and Stuart, 2009). The data also agree well with the early (Svoboda et al., 1996) and more recent (Bloodgood and Sabatini, 2005; Takasaki and Sabatini, 2014; Tønnesen et al., 2014) measurements, which showed low diffusional resistance of the spine neck corresponding to low R_neck_. Particularly solid experimental evidence in agreement with our findings is the measurement of the diffusional resistance of the spine neck to Na^+^ ion flux which mediates the rapid removal of sodium from the spine head (Miyazaki and Ross, 2017, 2022). Due to the close analogy between diffusional coupling and the electrical resistance, it is difficult to argue against strong implications of diffusional measurements for the upper bound of the possible electrical resistance of the spine neck. An additional argument in favor of low R_neck_ and against the electrical role of spines are two reports based on indirect, Ca^2+^-imaging data. These studies provided strong evidence for the lack of correlation between the morphology and dimensions of the spine neck and uncaging-evoked EPSPs and Ca^2+^ transients (Takasaki and Sabatini, 2014; Bywalez et al., 2015). Our results imply that spines on thin basal dendrites are characterized by uniform electrical behaviour regardless of considerable natural morphological variations (Jones and Powell, 1969; Tønnesen et al., 2014). Thus, the data argue that relatively small changes in the morphology of individual spines known to occur following induced synaptic plasticity (Takasaki et al., 2013; Araya et al., 2014; Tønnesen et al., 2014; Tazerart et al., 2020) are likely to be the by-product and not the cause of plastic changes.

The results supporting low Rneck call into question several existing hypotheses regarding the electrical role of spines. One group of studies based on indirect, Ca^2+^-measurements and numerical simulations postulated that spines neck filters EPSPs (Araya et al., 2006b, 2014), that spines cause the reduction of location-dependent variability of local EPSP amplitude and normalize synaptic activation of NMDA receptors and voltage-gated channels (Gulledge et al., 2012), that spines amplify EPSPs up to 45-fold thus facilitating electrical interactions among coactive inputs and promoting associated forms of plasticity and storage (Harnett et al., 2012), that spine neck resistance plays a vital role in determining the spine EPSP amplitude (Acker et al., 2016), that spine geometry plays a key role in shaping the EPSP time course (Cartailler et al., 2018; Lagache et al., 2019). It is certainly possible that some of these discrepancies are caused by the differences in the preparations and experimental methods and conditions. At the same time, the experimental evidence for the above hypotheses cannot be interpreted unambiguously because Ca^2+^ signals are an indirect, slow, and highly non-linear indicator of transmembrane voltage changes. Additionally, as a rule, the results of numerical simulations in neurophysiology depend on approximations and uncertain assumptions. Our recordings of electrical signals on two sides of the spine neck do not support the above hypotheses.

Another group of studies combined voltage imaging and numerical simulations to determine the electrical role of spines (Acker et al., 2016; Kwon et al., 2017; Cornejo et al., 2022). They interpreted their results as supporting high R_neck_ and a significant attenuation of EPSP signals in the parent dendrite. In an *in vivo* study using two-photon microscopy and a genetically encoded protein voltage probe Cornejo et al. (2022) concluded that spines are isolated voltage compartments that could be important for dendritic integration and disease states. However, none of those studies based on recording fluorescence light from voltage-sensitive probes had adequate sensitivity and spatiotemporal resolution to detect and reconstruct EPSP signals simultaneously from the spine head and the parent dendrite. Therefore, the interpretation of these results is uncertain. Our data do not support conclusions from these reports.

### Contribution of an individual synapse to electrical signaling

Using high-sensitivity voltage imaging, we obtained unique data on the temporal summation of unitary EPSPs at the site of origin -i.e., at single excitatory synapses on dendritic spines. Our study shows that: (1) Temporal summation of repetitive quantal EPSP inputs at the site of origin (spine head) is markedly sublinear; (2) AMPAR desensitization combined with the reduction of the synaptic driving force seems to be an important determinant of sublinear EPSP summation; (3) Distinctly sub-linear uEPSP summation at the synaptic site is paralleled by near-linear summation at the soma-axon region; (4) Repetitive activation of individual synapses does not initiate APs at the soma-axon region or a dendritic spike, independently of the input frequency. In conclusion, local individual uEPSPs have fast kinetics resulting in little or no temporal summation at frequencies below 100 Hz. At a higher frequency (200 Hz), unitary EPSPs summate locally in a pronounced sub-linear manner. At the soma-axon region, EPSPs at both low and high frequency summate in a near-linear fashion due to the broadening of the EPSP signal caused by RC filtering in the dendrite. Thus, our study shows that the sublinear summation of EPSP at the synaptic sites prevents depolarization buildup and synaptic saturation. This mechanism widens the dynamic range of near-linear summation of repetitive EPSPs at the soma-axon region.

## Acknowledgment

We thank Lawrence B Cohen (Yale University) for continuous support and Leslie M. Loew (Centre for Cell Analysis and Modelling, UConn Health Centre) for kindly providing voltage-sensitive dyes. We thank Valentina Emiliani (Institut de la Vision, Paris, France) and Dimitrii Tanese (Paris Descartes, Paris, France) for collaborating on setting up and using computer-generated holography. This work was supported by the National Institute of Health MH106906.

## Supplementary material

**Figure S1.**
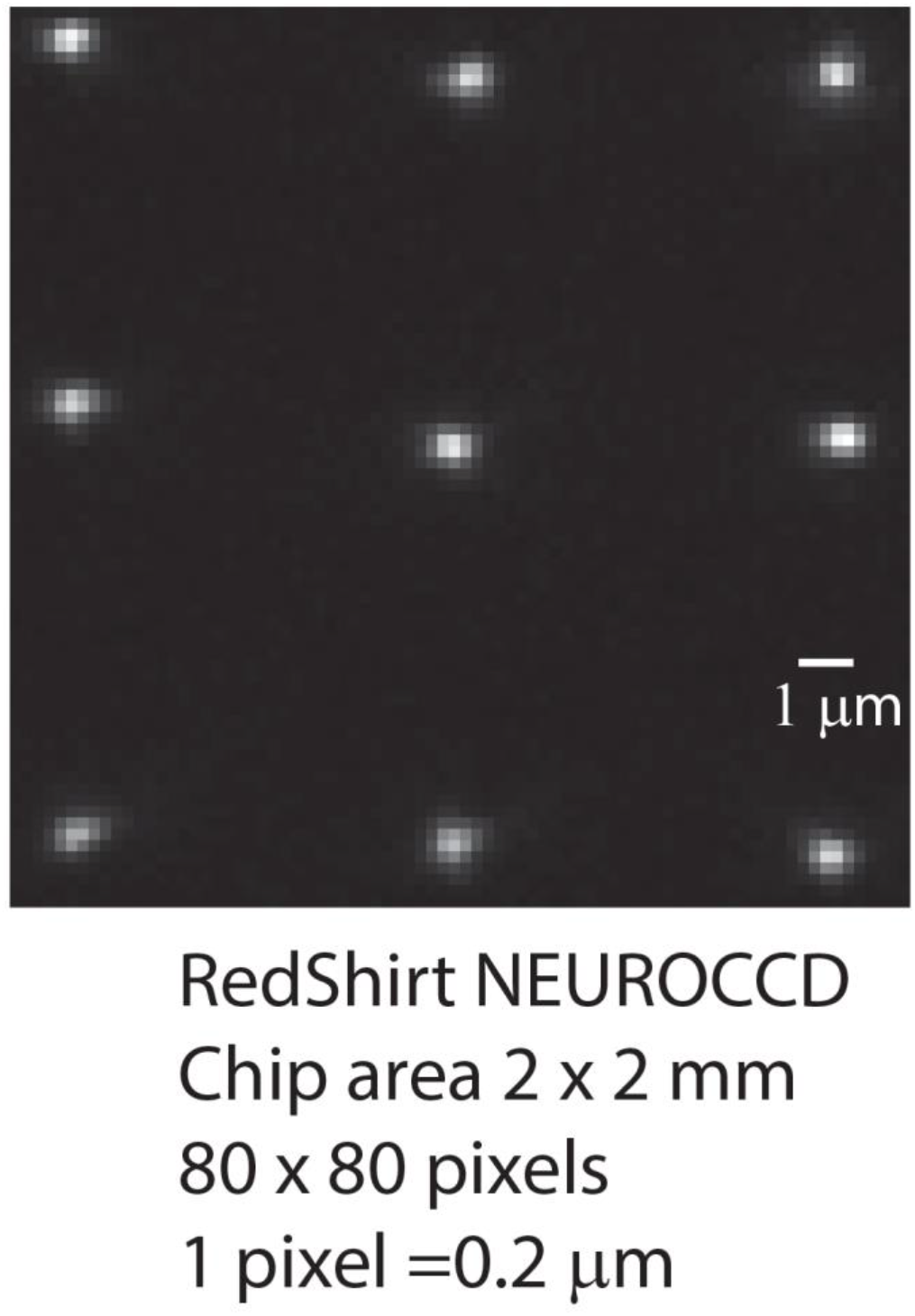
Patterned holographic illumination was recorded from a thin layer of rhodamine 6G dispersed on a microscope slide. The 720 nm spot covered ∼3 pixels corresponding to a size of ∼600 nm

**Figure S2.**
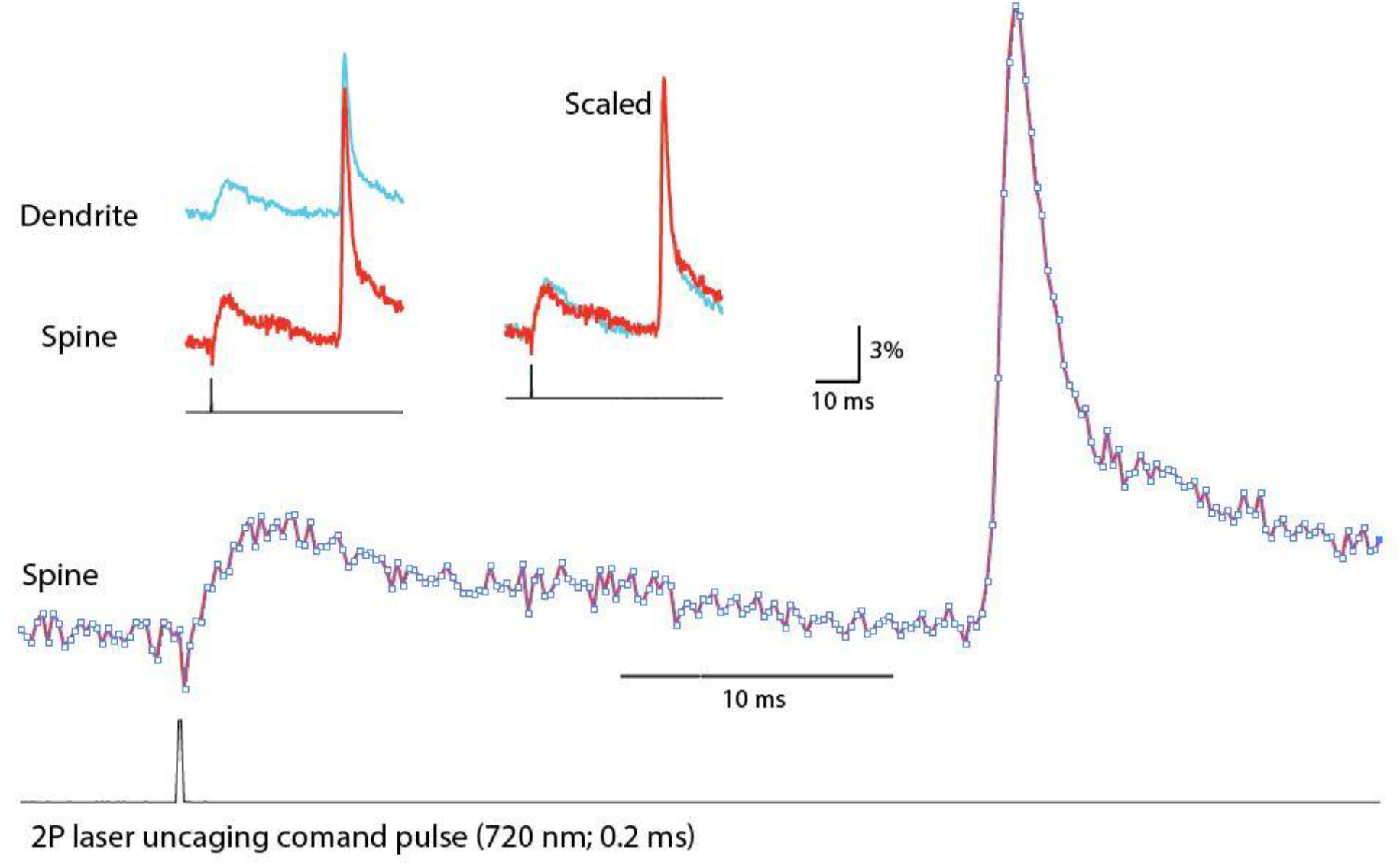
Stimulated Emission Depletion (STED) effect of the uncaging 720 nm red light reducing the intensity of voltage-sensitive die fluorescence exited by 532 nm excitation light. Upper traces: raw signals from dendrite and spine and superimposed scaled signals. Bottom black trace: uncaging command pulse. Lower traces: superimposed scaled signals on an expanded time scale. Bottom black trace: uncaging command pulse. The short (0.2 ms) uncaging pulse controlled by a Pockels cell modulated only one data point during the 5 kHz recording of the voltage-sensitive die signals. The peak amplitudes of the uEPSP and bAP were not influenced by the STED effect.

**Figure S3.**
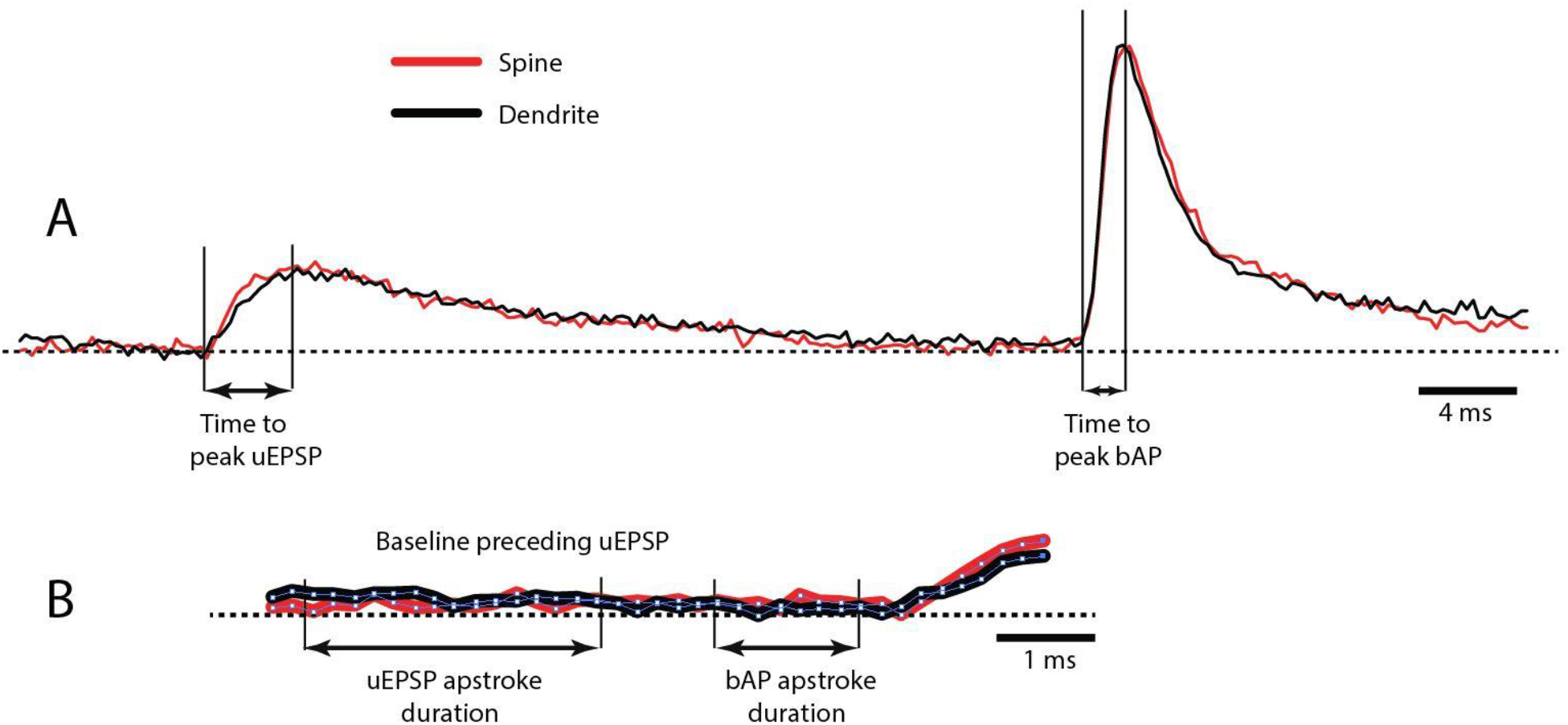
The absence of residual die bleaching effect on uEPSP and bAP signals shape and size. **A**, Optical recordings of uEPSP and bAP signals from the spine head and parent dendrite. Duration of uEPSP and bAP upstroke indicated by arrows. **B**, Baseline optical signals from the start of recording to the start of the uEPSP on an expanded time scale. Note that, during a period equivalent to the upstroke of the uEPSP and bAP, changes in signal amplitude due to the residual bleaching after bleaching correction (Methods) are smaller than the noise in recordings. We conclude that residual bleaching did not affect signal amplitudes.

